# Relative sensitivity of explicit re-aiming and implicit motor adaptation

**DOI:** 10.1101/308510

**Authors:** Sarah A. Hutter, Jordan A. Taylor

## Abstract

It has become increasingly clear that learning in visuomotor rotation tasks, which induce an angular mismatch between movements of the hand and visual feedback, largely results from the combined effort of two distinct processes: implicit motor adaptation and explicit re-aiming. However, it remains unclear how these two processes work together to produce trial-by-trial learning. Previous work has found that implicit motor adaptation operates automatically, regardless of task relevancy, and saturates for large errors. In contrast, little is known about the automaticity of explicit re-aiming and its sensitivity to error magnitude. Here we sought to characterize the automaticity and sensitivity function of these two processes to determine how they work together to facilitate performance in a visuomotor rotation task. We found that implicit adaptation scales relative to the visual error, but only for small perturbations – replicating prior work. In contrast, explicit re-aiming scales linearly for all tested perturbation sizes. Furthermore, the consistency of the perturbation appears to diminish both implicit adaptation and explicit re-aiming, but to different degrees. Whereas implicit adaptation always displayed a response to the error, explicit re-aiming was only engaged when errors displayed a minimal degree of consistency. This comports with the idea that implicit adaptation is obligatory and less flexible, while explicit re-aiming is volitional and flexible.

## Introduction

Compensating for movement errors is critical to the motor learning process. Thus, characterizing the sensitivity of the behavioral response to these errors should reveal fundamental principles and constraints of the motor system. For over twenty years, the motor control field has sought to characterize this sensitivity function using system identification techniques borrowed from engineering. These techniques generally consist of imposing a transient and often varying perturbation on the system to observe the behavioral response.

Despite employing this theory-driven and elegant approach, the observed sensitivity functions have been highly variable and appear to depend on a number of experimental factors. In a seminal study, Scheidt and colleagues (2001) found that the motor system adapted to transient and random force perturbations on a trial-by-trial basis in a force-field-adaptation task. This adaptive response appears to be sensitive to the direction of the perturbation, but insensitive to both the timing and magnitude (i.e., strength) of the perturbation (Fine and Thoroughman, 2006). However, if the perturbations are drawn from a non-zero mean distribution (Fine and Thoroughman, 2007), occur frequently (Fine and Thoroughman, 2007), or are applied in a consistent fashion (Castro et al., 2014), then the adaptive response becomes more sensitive.

Similar results have been found in studies of visuomotor rotations: The adaptive response appears to be highly sensitive to the direction of the rotation, but less sensitive to its magnitude (Butcher and Taylor, 2018). In fact, the time course of adaptation begins to saturate in response to rotations greater than ~ 6° (Wei and Kording 2010; Marko et al., 2012; Morehead et al., 2017; Kim et al., 2018), despite prolonged periods of training with both non-zero mean and consistent rotational perturbations (Morehead and Smith, 2017). This was made clear by employing a task-irrelevant error clamp task, in which cursor feedback of the supposed hand’s position is consistently offset from the target path by a fixed angular value regardless of the hand’s true location (Morehead et al., 2017). Despite the task-irrelevance of this feedback, robust adaptation is observed even when movement angles are uncorrelated with the angle of the rotational perturbation.

At first glance, the lack of sensitivity of this adaptive response is puzzling given the variety of motor behaviors and skills we can employ. However, when subjects have control of the angular position of the cursor – thus, making it task-relevant – sensitivity is restored (Morehead et al. 2017). This suggests that additional learning processes, such as explicit re-aiming, may play a role in making the appropriate behavioral response given the demands of the task (Heuer and Hegele, 2008; Taylor et al., 2014; Bond and Taylor, 2015). Indeed, the varying sensitivity functions measured in the aforementioned studies vacillated around conditions when the subjects could potentially affect the outcome in the task. For example, Wei and Körding (2009) used a task in which visual error changed randomly on every trial. In this situation, aiming anywhere except straight toward the target would actually be counter-productive. In contrast, the adaptive response become more proportional when force perturbations have a non-zero mean (Fine and Thoroughman 2007). Similarly, when force perturbations were made to be more consistent, by increasing the number of trials in a row for which a particular perturbation is present, the learning rate increased (Castro et al., 2014).

While we note that there are other substantial differences between these aforementioned studies, such as rotational versus force perturbations, we hypothesize that controllability over performance is a critical factor, which, in turn, engages explicit re-aiming to restore performance. Specifically, we propose that explicit re-aiming may only play a role when the perturbations are consistent and the visual errors are quite large (and/or implicit adaptation is saturated). Here, in two experiments, we set out to test these ideas by examining the sensitivity functions of both explicit re-aiming and implicit adaptation to a range of perturbations that varied in their degree of consistency.

## Materials and Methods

### Participants

Eighty human subjects [57 women, mean age 20.4 (SD 3.24) yrs.] were recruited for Experiment 1, and 26 subjects [18 women, mean age 19.69 (SD 1.35) yrs.] were recruited for Experiment 2. The sample size for Experiment 1 was guided by the typical convention of 10-20 subjects per condition (4 conditions) for visuomotor adaptation tasks. The sample size for Experiment 2 was determined by an *a priori* power analysis using the slope of explicit re-aiming sensitivity function from Experiment 1 between the Consistent-2 and Consistent-7 conditions. This analysis determined that we needed 13 subjects per condition to obtain robust power. All subjects were drawn from the research participation pool maintained by the Department of Psychology at Princeton University and received either course credit or monetary compensation for participating. Subjects were right handed, as verified by the Edinburgh handedness Inventory (Oldfield 1971), and self-reported having normal or corrected-to-normal vison. The experimental protocol was approved by the Princeton University Institutional Review Board and all subjects provided written, informed consent.

### Apparatus

Subjects preformed horizontal movements in a center-out reaching task. These movements were recorded with a digitizing pen and Wacom tablet, with the tablet sampling movement trajectories at 60 Hz. All stimuli were displayed by a 17-in., Planar touch sensitive monitor with a refresh rate of 60 Hz and computed by a Dell OptiPlex 7040 machine running Windows 7. The touch sensitive monitor allowed subjects to report their intended movement by simply tapping on the screen (Fig. 1; Bond and Taylor, 2017). Visual feedback of the hand was obstructed by the monitor, which was mounted 25 cm above the tablet. A small circular cursor (0.15-cm radius) provided feedback information to subjects. The game was controlled by custom software coded in Matlab Psychtoolbox (The Mathworks, Natick, MA).

**Figure 1.**
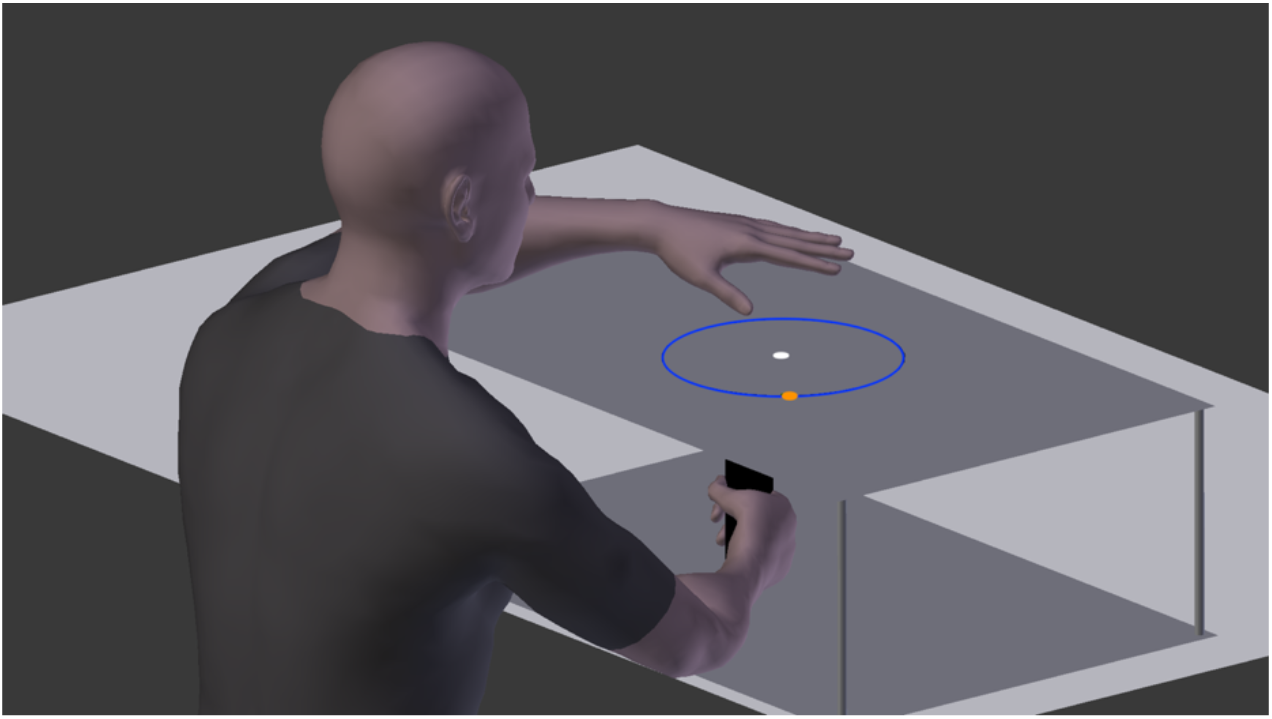
Experimental setup for both experiments. The subject holds a digitizing pen in his right hand, which is covered by the touch screen. The left hand is used to tap the touch screen on the blue ring to report aiming location before each trial. After the subject taps on the screen, the blue aiming ring would disappear and the target would turn green.

### Procedure

Subjects began each trial with their right hand at the center of the visual workspace. After holding this position for 500ms, a circular orange target (0.25 cm radius) appeared 7 cm from the starting position. Appearing along with the target, an “aiming” ring consisting of a blue circle that was centered on the starting location and had a radius of 7 cm (Fig. 1). Subjects were instructed to report their intended aiming location by tapping the aiming ring on the surface of the touch screen with their left hand. Once a touch was recorded, the target turned from orange to green, the aiming ring disappeared, and subjects were able to reach with their right hand. If a subject attempted to reach with their right hand prior to tapping the touchscreen with their left hand, a message “Remember to report aim” was displayed and the trial was restarted.

Subjects were instructed to make a fast, straight “shooting” movement through the target with their right hand. They were informed that it was not necessary to stop on the target, but that they should be careful to move far enough to pass completely through the target. Subjects were provided with continuous, online feedback of the cursor throughout the movement. Once the subject’s hand passed 7 cm, endpoint feedback was displayed for 1 s. If the final position of the cursor overlapped with the target, subjects heard a pleasant “ding” sound; otherwise, they heard an unpleasant “buzz.” If the time from leaving the start position to reaching out 7 cm exceeded 800 ms, the feedback “too slow” was given (this occurred on approximately 1% of trials, and these trials were excluded from further analysis). Following feedback presentation, subjects were guided back to the start position by a white ring that was centered on the starting location and whose radius represented the distance between subjects’ hand position and the starting location. Veridical feedback of the cursor was restored once the hand was within 1 cm of the starting position.

We pseudorandomized the target locations across the workspace and across subjects so that any potential visual or biomechanical biases would average out. On each trial, the target could appear in one of eight locations on the aiming ring. However, the angular configuration of the target locations differed across subjects. There were five sets of target configurations, where the ‘first’ target could be located at 0°, 9°, 18°, 24°, or 36° relative to the x-axis and the targets were always spaced 45° apart. Each subject was exposed to only one configuration of target locations.

To assay the sensitivity function of implicit adaptation and explicit re-aiming, an angular rotation of ± 0°, 2°, 4°, 8°, and 16° was imposed on the cursor in Experiment 1. These rotation sizes were chosen to correspond to the set of lateral displacements used in Wei and Körding (2009). Furthermore, the rotations were counterbalanced such that the mean rotation size over the experiment was 0°. For experiment 2, a 32° rotation was exchanged for the 2° rotation, such that the rotational perturbations imposed during the task were ± 0°, 4°, 8°, 16°, and 32°.

A secondary goal of this experiment was to determine if the sensitivity function changed based on the consistency of the rotation (i.e., how frequently the rotations changed during training), which has been reported by prior studies (Fine and Thoroughman, 2007; Castro et al., 2014). To this end, in Experiment 1 subjects were equally divided into four groups: Consistent-1, Consistent-2, Consistent-3, and Consistent-7. In the Consistent-1 condition, the rotation changed on every trial, effectively making this an inconsistent condition; though the target location remained the same for 7 trials. In the Consistent-2, Consistent-3 and Consistent-7 condition, each “mini-block” consisted of 2, 3 or 7 trials, respectively, where the rotation changed after each mini-block. For these conditions, the target location also changed at the onset of each mini-block. In all cases, visual perturbations were pseudo-randomly generated such that no rotation size was immediately repeated and each rotation size occurred at each target location at least once. Experiment 2 consisted of only the Consistent-2 and Consistent-7 conditions, which had mini-block lengths of 2 and 7 trials, respectively.

The experiments proceeded by first providing veridical feedback for eight familiarization trials and then pseudo-random visual rotations of the cursor for 504 trials in the Consistent-1 and Consistent-7 conditions and 432 trials in the Consistent-2 and Consistent-3 condition. The discrepancy in trial length is the result of complete counter-balancing. The entire experiment took approximately one hour.

Note, subjects were informed that the mapping between their hand and the cursor may change during the experiment and that they should tap on the aiming ring where they intend to aim in order to hit the target. Subjects were not told the nature of the visual perturbation nor when the visual disturbance would be in effect during the experiment.

### Data and statistical analyses

All data and statistical analyses were performed in Matlab. The digitizing tablet recorded the trajectory of the right hand and the touchscreen monitor recorded the terminal position of the location tapped with the left hand. These data were transformed to define heading hand angles and aiming angles during training as follows: The hand angle trajectories and aiming locations were transformed from Cartesian to polar coordinates and rotated to a common axis with the convention that the target was positioned at 0° (directly to the right). As our primary interest concerns only the feedforward portion of the reach, we focused on the initial heading of the hand angle by examining the average angle of the hand between 1 and 3 cm into movement. Aiming angle was defined as the angle between the target and the tapped location on the touchscreen. Implicit adaptation was calculated by subtracting the subjects’ aiming angle from their hand angle (Taylor et al., 2014; Bond and Taylor, 2017). For all measures, positive angles indicate a counterclockwise deviation from the target. As we have only two measured values, with implicit adaptation being computed as the subtraction of aiming angle from hand angle, we performed statistical analyses only on aiming angles and implicit adaptation angles – our primary variables of interest – and we report hand angles only for completeness.

In the following statistical analyses, unless otherwise specified, only the second trial of each block was used. This allowed us to control for the confounding additive effects inherent in having different length mini-blocks for each condition. For predictive purposes, the rotation size is considered to be the rotation size of the mini-block. Thus, subjects experienced the rotation on the first trial of the mini-block and we evaluated their response on the next (second) trial. For the Consistent-1 condition, the rotation size is considered to be the rotation experienced on the previous trial (n-1, where n is the trial being evaluated).

To quantify the sensitivity function for each consistency condition, we fit separate linear functions to each subject’s aiming angles and implicit adaptation angles with respect to the rotation size. A significant slope indicates that the subject changed their behavior in response to the error. Differences in slopes between consistency conditions were evaluated by submitting the slopes to a one-way ANOVA; post-hoc t-tests were conducted when appropriate and corrected using the Bonferroni method. We also sought to determine if the overall slope of the sensitivity function was similar across rotation sizes. Previous studies have reported that the sensitivity of the response scales with the rotation size before reaching a saturation point at higher rotation magnitudes (Wei and Kording 2009; Morehead et al., 2017). To assess this possibility, we adopted the method of Wei and Kording (2009) where they fit a second linear function to the range from −4 to 4°, which corresponds to lateral displacement between −2 and 2 cm in their study. The slopes of this second function and the overall function were compared with a pairwise t-test.

The intercept (or offset) of these functions is of less interest, but significant intercepts could be viewed as an accumulation of learning throughout training or the development of a more general bias during training (Ghilardi et al. 1995).

## Results

### Experiment 1

In this experiment, we sought to assess the sensitivity of implicit adaptation and explicit re-aiming as a function of the magnitude and consistency of rotational perturbations, which ranged from 0-16° within a subject and changed every 1, 2, 3, or 7 trials across subjects. Subjects attempted to counteract these perturbations for all consistency conditions, as can be seen by the change in the angle of the hand in response to the imposed error (Fig. 2A). For the majority of conditions, these changes in hand angle are the result of the combined output of implicit adaptation (Fig. 2B) and explicit re-aiming processes (Fig. 2C). To quantify the sensitivity of these processes, we fit a linear function to each process for each subject over the imposed rotations. For all consistency conditions, we found that the slope of the sensitivity function was significant for both implicit adaptation (p < 0.01) and explicit re-aiming (p < 0.001; see Table 1). The intercept of these linear fits was not different from zero, except in the case of Consisitent-1 which had a small yet significant positive shift for both implicit adaptation (mean = 0.209°; t(19) = 2.573, p = 0.02) and explicit re-aiming (mean = 0.29°, t(19) = 3.218, p = 0.005), suggesting that learning largely did not accumulate throughout the experiment.

**Figure 2.**
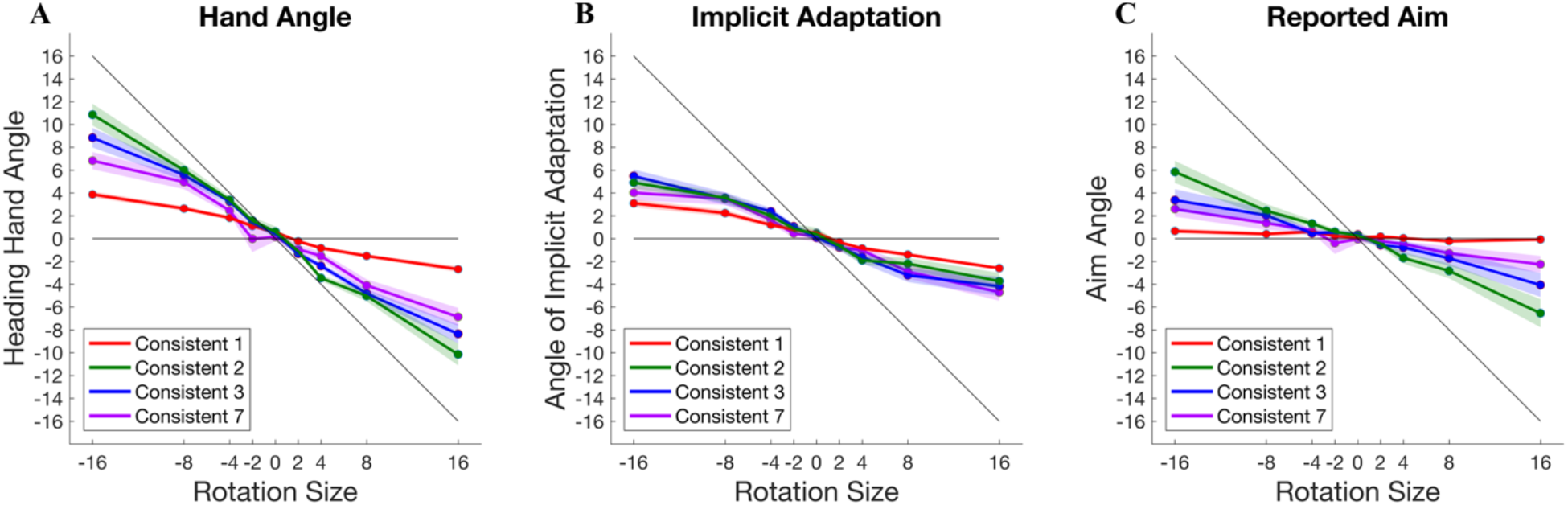
Responses to imposed visual perturbations of −16 to 16° for each consistency condition: Consistent-1 (red), Consisten-2 (green), Consistent-3 (blue), and Consistent (purple). A) Angle of the hand heading direction. B) Angle of calculated implicit adaptation (hand angle – explicit re-aiming). C) Angle of explicit re-aiming, which was reported by touching screen with left hand. Shaded regions represent standard error.

**Table 1.** Statistics for the slope of the linear fit between imposed rotation and response: implicit adaptation or explicit re-aiming. The linear equation was fit to the data of each subject.

| **Implicit Adaptation - Slope of Linear Fit** |  |  |  |  |
| --- | --- | --- | --- | --- |
|  | Mean | Confidence interval | Mean r-value | Percent of p-values < 0.05 |
| Consistent 1 | -0.192 | -0.214 to -0.170 | 0.894 | 95.0% |
| Consistent 2 | -0.298 | -0.365 to -0.231 | 0.824 | 90.0% |
| Consistent 3 | -0.335 | -0.423 to -0.248 | 0.869 | 95.0% |
| Consistent 7 | -0.300 | -0.363 to -0.237 | 0.789 | 80.0% |

| **Explicit Re-Aiming - Slope of Linear Fit** |  |  |  |  |
| --- | --- | --- | --- | --- |
|  | Mean | Confidence interval | Mean r-value | Percent of p-values < 0.05 |
| Consistent 1 | -0.028 | -0.045 to -0.012 | 0.488 | 25.0% |
| Consistent 2 | -0.374 | -0.508 to -0.240 | 0.794 | 75.0% |
| Consistent 3 | -0.230 | -0.352 to -0.108 | 0.702 | 60.0% |
| Consistent 7 | -0.152 | -0.232 to -0.071 | 0.651 | 55.0% |

Despite finding significant slopes for all the implicit adaptation sensitivity functions, it can be seen in Figure 2B, that they are do not appear to be perfectly linear: The sensitivity appears to saturate for large rotations, which has been observed in previous studies (Wei and Kording 2009; Morehead & Smith 2017). To determine if implicit adaptation sensitivity is better described as a piecewise function, we followed the method of Wei and Kording (2009) to compare the slope for small rotations versus the overall function. We find that these slopes are different for the Consistent-1 (t(19) = 2.971, p = 0.008), Consistent-2 (t(19) = 3.227, p = 0.004), and Consistent-3 conditions (t(19) = 2.941, p = 0.008). Note, the Consistent-1 condition is nearly identical to the study by Wei and Kording (2009), replicating their findings. However, the slopes were not different for the Consistent-7 condition. Given the visual similarity in the functions between consistency conditions, we suspect that this is likely attributable to noise – an issue we will address in Experiment 2. We performed the same comparisons between slopes within each consistency condition for explicit re-aiming and found no significant differences (all ps > 0.05), although it appears that there may be differences in the slopes between consistency conditions.

To determine if there were significant differences in the slope of the sensitivity function between consistency conditions, we submitted the slopes for both implicit adaptation and explicit re-aiming to separate one-way ANOVAs. For implicit adaptation, while we find a significant difference between conditions (F(3) = 3.55, p-value = 0.02), this effect is driven by the difference between the Consistent-1 condition and all other conditions: Consistent-2 (t(38) = 2.958, p-value = 0.005), Consistent-3 (t(38) = 3.099, p-value = 0.003), Consistent-7 (t(38) = 3.166, p-value = 0.003). This suggests an overall reduction in the sensitivity of implicit adaptation when there is no consistency in error (Fig. 2B). Explicit re-aiming also differs with consistency (F(3) = 8.15, p < 0.001). Interestingly, post-hoc t-tests reveal that increasing the consistency by a single trial radically increases the sensitivity, compare Consistent-1 and Consistent-2 conditions (t(38) = 5.02, p < 0.001, Fig. 2C). However, as the consistency is further increased the sensitivity function tends to decrease, comparing Consistent-2 with Consistent-3 (t(38) = 1.561, p = 0.13), Consistent-2 with Consistent-7 (t(38) = 2.789, p = 0.008), and Consistent-3 with Consistent-7 (t(38) = 1.05, p = 0.30, Fig. 2C).

We compared the slope of implicit adaptation and explicit re-aiming within each condition to determine if one was more sensitive to the imposed perturbation. Implicit adaptation was significantly more sensitive to perturbation size than explicit re-aiming in the Consistent-1 condition (t(19) = 11.620, p < 0.001) and in the Consistent-7 condition (t(19) = 2.433, p = 0.02). Greater sensitivity of implicit adaptation over explicit re-aiming in the Consistent-1 condition suggests that explicit re-aiming can be “turned off” when it is not useful, while implicit adaptation proceeds regardless. There was no difference in the slopes of implicit adaptation and explicit re-aiming in the Consistent-2 (t(19) = 0.810, p = 0.43) and Consistent-3 conditions (t(19) = 1.078, p = 0.30). The relative magnitude of explicit re-aiming and implicit adaptation are generally equivalent when learning is useful to task performance.

It is worth noting that our analysis of sensitivity focused only on the changes in behavior following the first experience with a new rotation in a mini-block. For the Consistent-2, −3, and - 7 conditions, the same rotational perturbation continued for additional trials. Thus, somewhat trivially, subjects could continue to implicitly adapt and explicitly re-aim. This is apparent in Figure 3, although the response appears to negatively accelerate with continued training in the mini-block, which is likely attributable to progressively decreasing visual errors. Consequently, these trials become increasingly contaminated by prior performance and thus provide an impure measure of the error sensitivity function. Therefore, we limited our error sensitivity function estimations to only the second trial of the mini-block.

**Figure 3.**
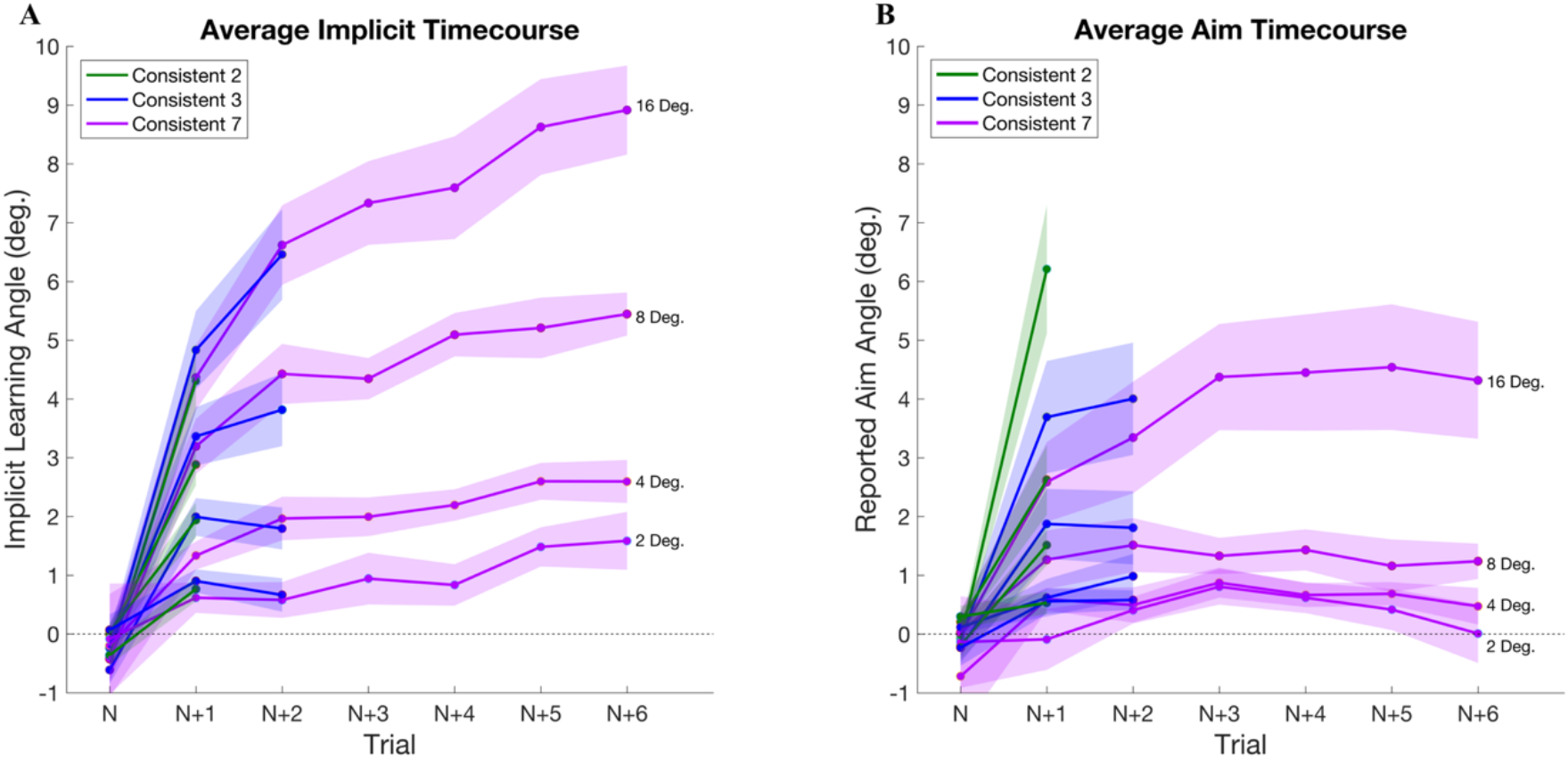
Time course of A) implicit adaptation and B) explicit re-aiming throughout each mini-block. Each mini-block is the average of the trials in a row for which each rotation size was consistent (2 trials for Consistent-2, 3 trials for Consistent-3, 7 trials for Consistent-7). Shaded regions represent standard error.

Although not considered in our statistical analyses, Figure 3A does highlight the degree of sensitivity that implicit adaptation shows for rotation magnitude. This can be seen in the separation of the time course for each rotation magnitude. There is even clear differentiation between rotations as close as 2° and 4° (Kim et al., 2018). Re-aiming, however, does not show as clean a separation (Fig. 3B).

Next, we were interested in determining if the change in explicit re-aiming sensitivity as a function of consistency was due to a fundamental feature of the learning process or as a result of a statistical property of the training environment. One possible explanation is that explicit re-aiming is actually sensitive to the magnitude of changes in apparent visual error between trials. For example, in the Consistent-7 condition, visual error was progressively smaller within a miniblock, but quite large in-between mini-blocks. In contrast, in the Consistent-1 condition, every trial was effectively in-between mini-blocks and, thus, larger visual errors were experienced more frequently. Indeed, the cumulative distribution of visual errors (CDF) varied significantly by condition for both means (F(3) = 87.98, p < 0.001) and standard deviations (F(3) = 8.11, p < 0.001, Fig. 4B). Most notably, the mean of the CDF for Consistent-2 is larger than the means the of Consistent-3 (t(38) = 3.032, p = 0.004) and Consistent-7 (t(38) = 10.2, p < 0.001). The standard deviation of the CDF for Consistent-2 condition was also larger than that of Consistent-3 (t(38) = 2.251, p = 0.03), and Consistent-7 (t(38) = 3.348, p = 0.002). These cumulative distributions suggest that decreasing the consistency of the perturbation increased the average change in visual error. This may account for the increase in the magnitude of the aiming response, with the caveat that this relationship breaks-down when the visual error is completely unpredictable. It should be noted that we cannot produce a causal claim with this experimental setup as changes in visual error are by definition influenced by aiming behavior.

**Figure 4.**
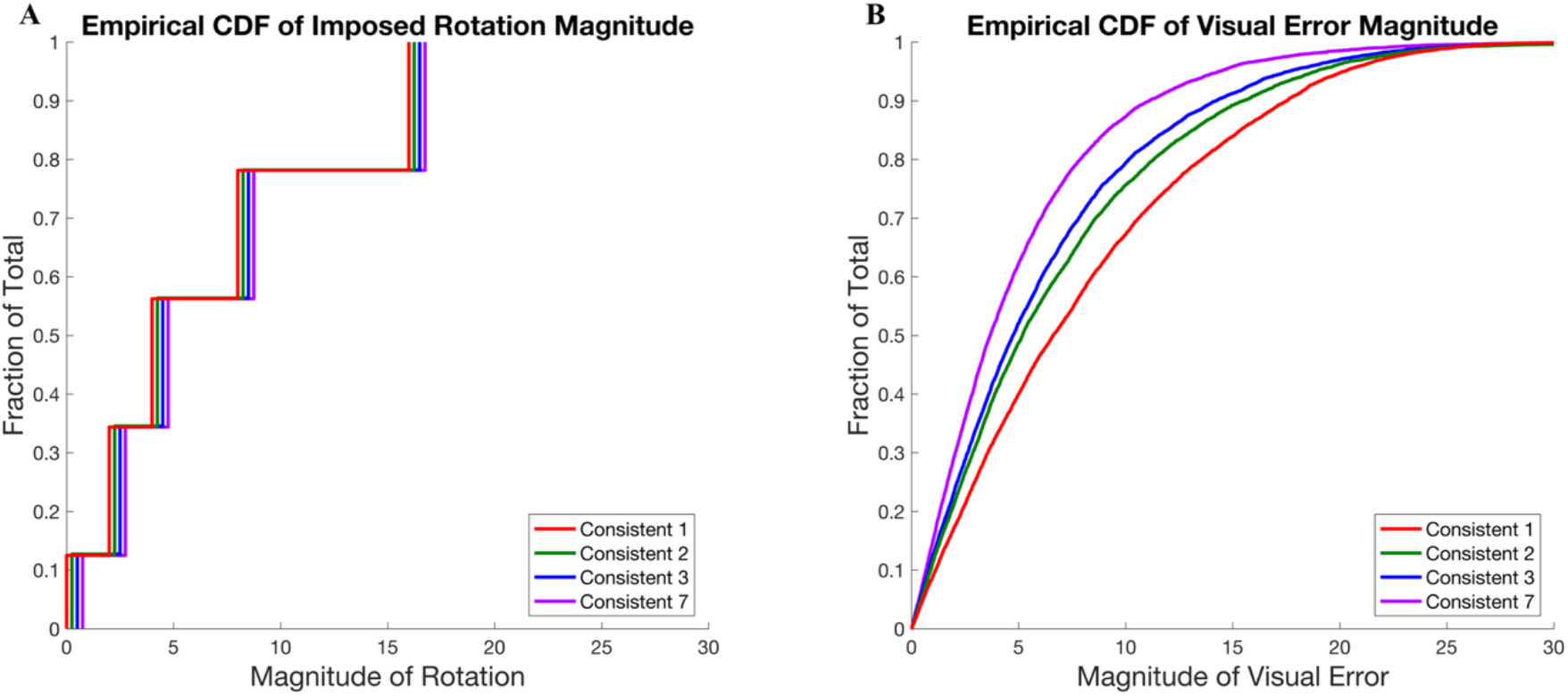
Empirical cumulative distribution function for A) imposed rotation magnitude and B) visual error magnitude. The imposed rotations are jittered for clarity.

In sum, we found that error sensitivity of implicit adaptation was largely invariant across consistency conditions. These findings are consistent with previous studies employing procedures to isolate implicit adaptation, although using prolonged block designs (Bond and Taylor, 2015; Morehead et al., 2017). Furthermore, we found that sensitivity functions for three out of the four consistency conditions tended to saturate for implicit adaptation, which is largely consistent with results from previous studies (Wei and Kording 2009; Morehead et al., 2017; Kim et al., 2018). However, the sensitivity function for implicit adaptation at the most consistent condition did not significantly saturate. In contrast, the sensitivity of explicit re-aiming changed as a function of consistency but showed linearity as a function of rotation size. To clarify these issues, we conducted a follow-up study (Experiment 2) to both replicate our central findings and extend them by including larger rotation sizes. Here we limited the study to two consistency conditions: Consistent-2 and Consistent-7.

### Experiment 2 Results

Similar to Experiment 1, implicit adaptation and explicit re-aiming were sensitive to the rotational perturbations for both consistency conditions. The average slope of implicit adaptation was significant for the Consistent 2 (t(12) = 7.389, p < 0.001) and Consistent-7 (t(12) = 3.827, p = 0.002) conditions (Fig. 5 and Table 2). Likewise, the slopes were significant for explicit re-aiming in the Consistent-2 (t(12) = 6.907, p < 0.001) and Consistent-7 conditions (t(12) = 3.685, p = 0.003). This replicates the sensitivity of implicit adaptation and explicit re-aiming to rotation size from the first experiment.

**Figure 5.**
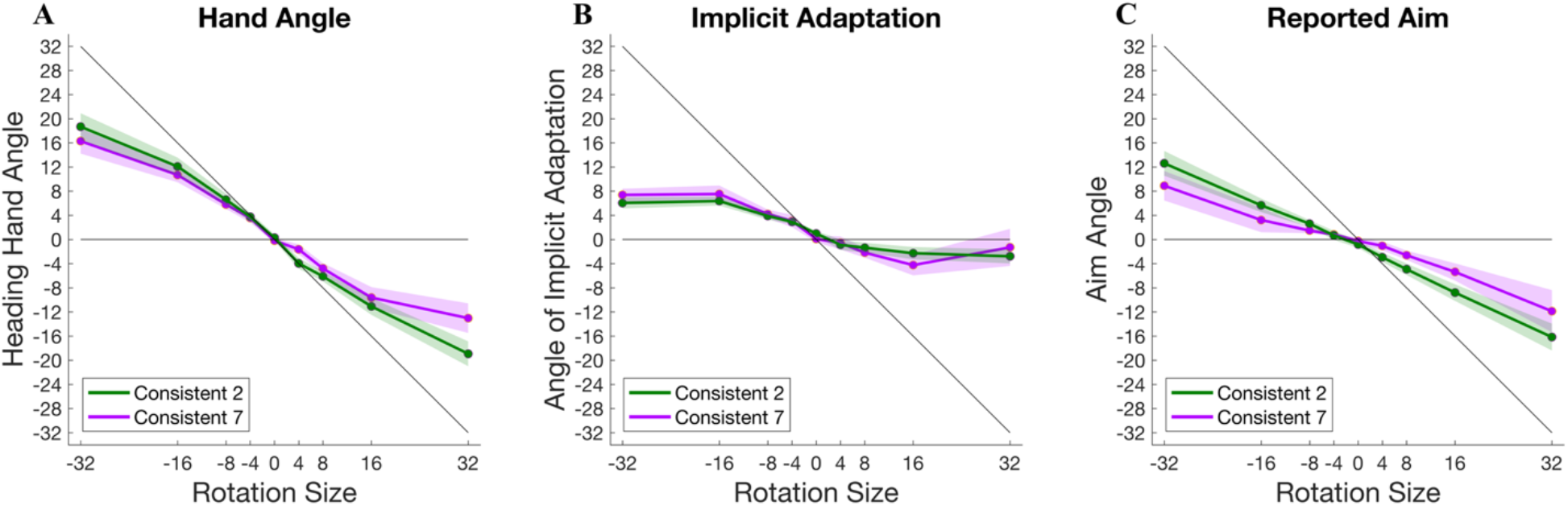
Responses to imposed visual perturbations of −32 to 32°. A) Angle of the hand heading direction. B) Angle of calculated implicit adaptation (hand angle – explicit re-aiming). C) Angle of explicit re-aiming. Shaded regions represent standard error.

**Table 2.** Statistics for the slope of the linear fit between imposed rotation and response: implicit adaptation or explicit re-aiming. The linear equation was fit to the data of each subject.

| Implicit Adaptation - Slope of Linear Fit |  |  |  |  |
| --- | --- | --- | --- | --- |
|  | Mean | Confidence interval | Mean r-value | Percent of p-values < 0.05 |
| Consistent 2 | -0.176 | -0.223 to -0.129 | 0.782 | 84.6% |
| Consistent 7 | -0.196 | -0.296 to -0.096 | 0.750 | 61.5% |

| Explicit Re-Aiming - Slope of Linear Fit |  |  |  |  |
| --- | --- | --- | --- | --- |
|  | Mean | Confidence interval | Mean r-value | Percent of p-values < 0.05 |
| Consistent 2 | -0.451 | -0.579 to -0.323 | 0.910 | 92.3% |
| Consistent 7 | -0.310 | -0.475 to -0.145 | 0.692 | 69.2% |

Unlike in Experiment 1, the intercepts of the linear fits to implicit adaptation were significantly larger than zero for both Consistent-2 and Consistent-7 conditions, although quite small (mean = 1.436° and 1.552° respectively). The intercept of the linear fits to explicit re-aiming were significantly below zero for the Consistent-2 condition (mean = −1.342°, t(12) = 2.881, p = 0.01) but not the Consistent-7 condition. These results suggest that the accumulation of learning or a bias did not strongly develop over training.

The non-linearity of implicit adaptation replicated in the Consistent-2 condition. Non-linearity was found when comparing the average slope between rotations of ±4° and the full range (±32°; t(12) = 3.124, p = 0.008), as well as when comparing between ±8° and the full range (t(12) = 3.886, p = 0.002). As in Experiment 1, the Consistent-7 condition did not show this effect for the comparison of the partial range ±4° to the full range (t(12) = 1.815, p = 0.09). However, the range from ±8° did have a significantly different average slope from the full range (t(12) = 3.878, p = 0.002). The same tests performed on explicit re-aiming produced no significant results, consistent with the findings from Experiment 1: Implicit adaptation shows a strong tendency to saturate at relatively larger perturbation sizes, while explicit re-aiming continues to contribute proportionately to learning throughout the whole range.

To compare sensitivity as a function of consistency, we submitted the slopes of implicit adaptation and explicit re-aiming to separate two-sample t-tests. As in Experiment 1, we found that the sensitivity of implicit adaptation did not change as a function of consistency (t(24) = 0.350, p = 0.73). Unlike Experiment 1, however, explicit re-aiming behavior between Consistent-2 and Consistent-7 was not significant (t(24) = 1.327, p = 0.20). This suggests that the subtle scaling of sensitivity of implicit adaptation and explicit re-aiming as a function of consistency, which we observed in Experiment 1, is not a robust effect.

A within-condition comparison between implicit adaptation and re-aiming showed that explicit re-aiming was more sensitive to these large perturbations than was implicit adaptation in the Consisitent-2 condition (t(12) = 3.844, p = 0.002). However, this did not hold for the Consistent-7 condition (t(12) = 0.932, p = 0.37). As in the first experiment, we do not see a reliable pattern indicating that implicit adaptation is more or less sensitive than explicit re-aiming when adaptive learning is relevant to task performance.

Finally, in Figure 6A we can see that implicit adaptation saturates to the larger perturbation sizes used in this experiment. This is in stark contrast to what was seen for the smaller perturbations in the previous experiment (Fig. 3A). Explicit re-aiming, on the other hand, does not saturate and shows a clean differentiation in the time courses for large perturbation sizes (Fig. 6B).

**Figure 6.**
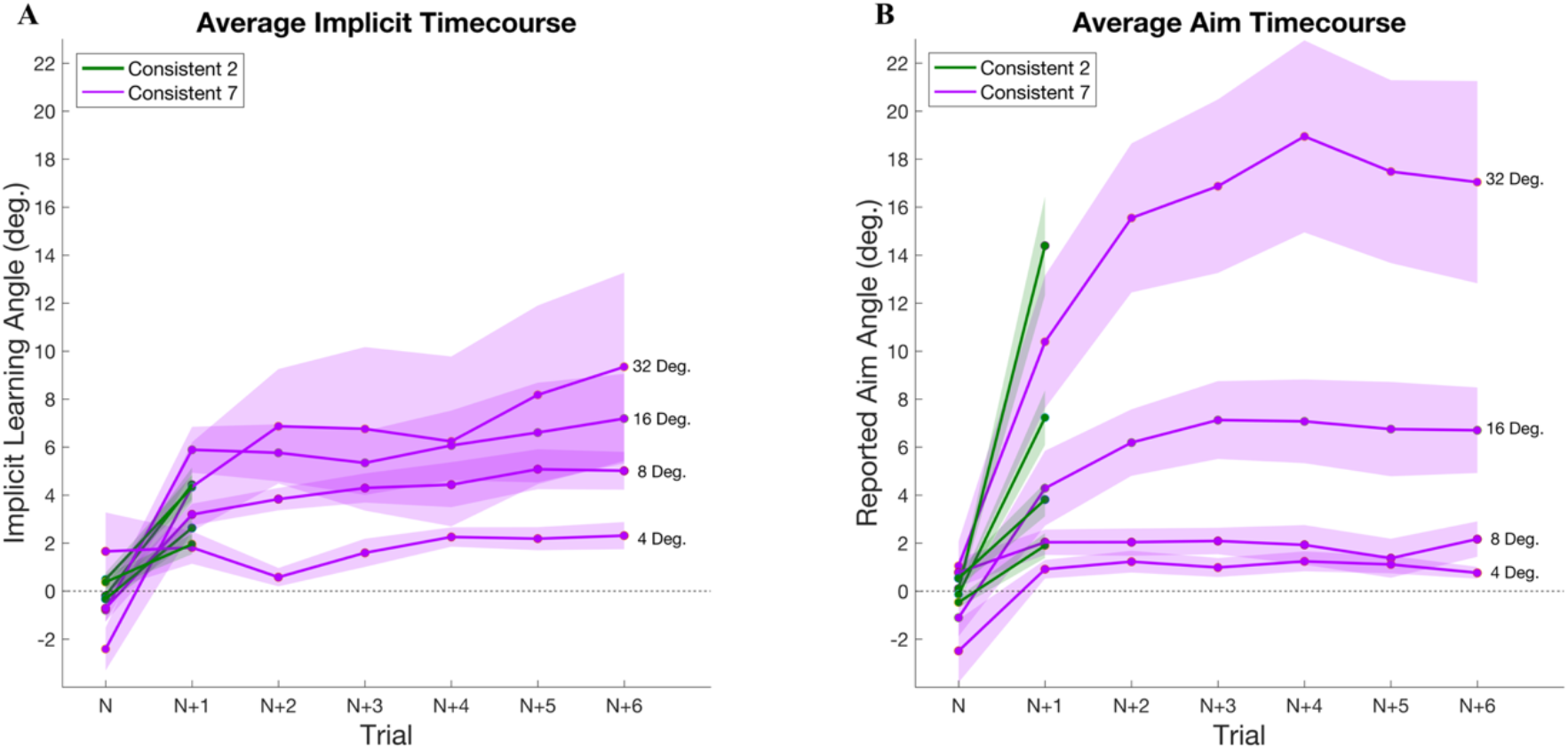
Time course of A) implicit adaptation and B) explicit re-aiming throughout each mini-block. Shaded regions represent standard error.

## Discussion

To determine the sensitivity function of implicit adaptation and explicit re-aiming we probed the motor system with small visual perturbations. In addition, we investigated the impact of error consistency on the aforementioned sensitivity of implicit and explicit processes by manipulating the number of trials in a row for which the perturbation was consistent. In two experiments, we perturbed visual feedback during center-out reaching movements. Both explicit re-aiming and implicit adaptation are sensitive to and respond differentially depending on the size the visual errors. By varying the consistency of the perturbation, we found that the sensitivity of implicit adaptation to small visual errors is impeded when the environment is completely unpredictable but stereotyped over all other levels of consistency. Likewise, the sensitivity of explicit re-aiming was practically null when the environment was inconsistent but stabilized with increased consistency in the perturbations. These results suggest that both implicit adaptation and explicit re-aiming are sensitive to very small perturbations, although implicit adaptation saturates for larger perturbations, and only minimal consistency of the perturbations is needed to stabilize these sensitivity functions.

Implicit adaptation was most sensitive to changes in error at very small error magnitudes and saturated between eight and sixteen degrees, in keeping with previous results (Wei and Kording 2009, Marko et al., 2012; Morehead et al., 2017; Kim et al., 2018). This non-linearity of implicit adaptation largely replicated across both experiments in most consistency conditions, with the only exception being the Consistent-7 condition. We suspect that the failure of this condition to replicate is likely the result of sampling error, as the general shape of its sensitivity curve is similar to those of the other conditions. Unlike implicit adaptation, explicit re-aiming showed greater sensitivity to large errors and produced a smaller differential response to changes in small errors (<8 deg.). These findings are consistent with previous studies suggesting that implicit adaptation is much more rigid than explicit re-aiming (Bond and Taylor 2015, 2017).

We hypothesized that previous accounts of environmental consistency influencing motor learning could be fully attributable to explicit processes. However, we found that implicit adaptation was also affected by consistency. Changing error magnitude on every trial produces less implicit adaptation than when there is some consistency to allow for predictive responses. This suggests that there is a minimum amount of error consistency necessary to achieve the full possible amount of implicit adaptation. Once this minimum consistency requirement is met, however, additional consistency does not produce greater adaptation. This finding modifies previous work suggesting that implicit adaptation is extremely stereotyped and insensitive to environmental features or task demands (Morehead et al., 2017).

Consistency was important to the magnitude of the re-aiming response. Moving from no error consistency to two trials in a row with the same error magnitude produced a sharp increase in explicit re-aiming. Surprisingly, further increases in error consistency decreased this response, as found in Experiment 1 (a similar trend was seen in Experiment 2). We believe that this reduction in sensitivity with increasing error consistency is the result of the statistical properties of our task: While the mean of the errors and the mean change in error from one trial to the next was zero for all conditions, the relative size of the change in error was larger for conditions with lower environmental consistency (see Fig. 4). This resulted in more circumstances where the error magnitude was very high in conditions with low error consistency as compared to those with high error consistency. Assuming, as previously argued, that the explicit re-aiming is more engaged when performance suddenly changes this would explain the modest increase in explicit re-aiming in low-consistency conditions.

Although not tested in these experiments, the sensitivity function of explicit re-aiming may be related to possible reward in the environment. Recent work has shown that explicit re-aiming is sensitive to bivalent feedback (Holland et al., 2018). One reason that explicit re-aiming is seemingly not sensitive to very small perturbations could be that the system categorizes these errors as “close enough” and treats them as correct (i.e. within the subject’s natural motor noise), allowing implicit adaptation to “clean up” the residual error. Under this framework, explicit re-aiming would be expected to begin making adjustments when movement is perceived to be sufficiently off-target. This phenomenon would likely be different across individuals and contexts.

The trade-off between implicit adaptation, which is most sensitive at small error sizes, and explicit re-aiming, which is most sensitive to large errors, suggest that the two may be fundamentally linked. Particularly, implicit adaptation allows for small updates to the internal model, necessary to avoid accumulation of small errors and to avoid drift in the system, while the explicit system can take over for fast learning of large perturbations. Whereas slow, gradual adaptation is sufficient for most learned tasks, sometimes a leap in learning is necessary to accommodate a radical environmental change. Whether implicit adaptation and explicit re-aiming are independent of one another, or if one relies on feedback from the other to successfully function, is an important question for future research.

Given mounting evidence, in this paper and others, that implicit adaptation saturates around the point at which explicit re-aiming is most sensitive suggests that the main function of implicit adaptation is calibrating to small errors. How then can we accomplish the full remapping necessary for skill learning? This re-mapping has been experimentally shown to be possible: Long-term adaptation studies conducted with subjects practicing throwing movements while wearing prism goggles showed that after weeks of practice subjects are able to make similarly precise movements immediately after donning the disrupting prism goggles as when they are not wearing them (Martin et al. 1996). One possibility is that implicit adaptation does eventually constitute the fully remapped motor behavior, but that this process proceeds on a far longer time-scale than usually studied in visuomotor adaptation experiments. However, recent work examining the time series of implicit adaptation under error-clamp conditions does not show the continuous, if slow, rise expected if implicit adaptation were to be eventually fully responsible for learning (Morehead et al., 2017). An extended, multi-day examination of explicit and implicit processes may provide insight into this open question.

## Acknowledgements

This work was supported by the National Institute of Neurological Disorders and Stroke (Grant R01 NS-084948 to J. A. Taylor)

